# AtPQT11, a P450 enzyme, detoxifies paraquat via N-demethylation

**DOI:** 10.1101/2021.06.23.449549

**Authors:** Yi-Jie Huang, Yue-Ping Huang, Jin-Qiu Xia, Zhou-Ping Fu, Yi-Fan Chen, Yi-Peng Huang, Aimin Ma, Wen-Tao Hou, Yu-Xing Chen, Xiaoquan Qi, Li-Ping Gao, Cheng-Bin Xiang

## Abstract

Paraquat is one of the most widely used nonselective herbicides in agriculture. Due to its wide use, paraquat resistant weeds have emerged and is becoming a potential threat to agriculture. The molecular mechanisms of paraquat resistance in weeds remain largely unknown. Physiological studies indicated that the impaired translocation of paraquat and enhanced antioxidation could improve paraquat resistance in plants. However, the detoxification of paraquat via active metabolism by plants has not been reported to date. Here we report that an activated expression of *At1g01600* encoding the P450 protein CYP86A4 confers paraquat resistance as revealed by the gain-of-function mutant *paraquat tolerance 11D* (*pqt11D*), in which a T-DNA with four 35S enhancers was inserted at 1646 bp upstream the ATG of *At1g01600*. The paraquat resistance can be recapitulated in Arabidopsis wild type by overexpressing *AtPQT11* (*At1g01600*), while its knockout mutant is hypersensitive to paraquat. Moreover, *AtPQT11* also confers paraquat resistance in *E. coli* when overexpressed. We further demonstrate that AtPQT11 has P450 enzyme activity that converts paraquat to N-demethyl paraquat nontoxic to Arabidopsis, therefore detoxifying paraquat in plants. Taken together, our results unequivocally demonstrate that AtPQT11/ CYP86A4 detoxifies paraquat via active metabolism, thus revealing a novel molecular mechanism of paraquat resistance in plants and providing a means potentially enabling crops to resist paraquat.

## Introduction

Paraquat (1,1’-dimethyl-4,4’-bipyridynium dichloride) has been widely used as a broad spectrum nonselective herbicide in agriculture for decades ^1^. Plants absorb paraquat from their environment and transport to chloroplasts, where it competes for electrons on the PSI and generates superoxide that is converted to H_2_O_2_ by superoxide dismutases (SODs), resultantly accumulating a large amount of ROS, which may lead to cell death ^2^.

Paraquat resistant weeds have emerged after the wide use of paraquat in agriculture. The underlying mechanisms of paraquat resistance in weeds have been studied mostly at physiological and biochemical levels ^3–6^. From these studies, two major mechanisms of paraquat resistance were suggested, including defects in paraquat translocation restricting the transport of paraquat to chloroplasts ^4,7^, and enhanced antioxidation scavenging ROS more efficiently ^8,9^. However, no genetic or molecular evidence for the proposed mechanisms has been revealed in weeds. With the power of the model plant Arabidopsis, several research groups have isolated paraquat resistant mutants, originally aiming at identifying novel components in oxidative stress response or cell death ^10,11^. These studies with Arabidopsis paraquat resistant mutants helped reveal the molecular mechanisms of paraquat resistance proposed by weed scientists based on their studies on paraquat resistant weeds. The unraveled molecular mechanisms from these studies fall into two categories; one involves paraquat uptake and transport ^12–15^, and the other is the enhanced ability scavenging ROS ^10,11,16,17^.

However, no evidence of paraquat degradation by plant active metabolism has been reported. Herbicide degradation by plants via active metabolism is a desirable mechanism for engineering crop resistance to herbicides. CYP450s (cytochrome P450s) constitute the biggest plant protein family and can mediate redox reaction in plant. They usually function as demethylase and/or hydroxylase in herbicide metabolism ^18^. Some P450s metabolize herbicides to lower toxic products which are transferred to vacuole as glucosyl conjugates ^19^. Two plant cytochrome P450 enzymes, CYP71A10 from *Helianthus tuberosus* and CYP76B1 from soybean, were reported having urea herbicide metabolism activity ^20^ In addition, tobacco CYP81B2 and CYP71A11 catabolize chlortoluron by ring methyl hydroxylation and N–methylation ^21^.

We previously isolated a number of paraquat tolerant Arabidopsis mutants and reported *pqt24* and *pqt3* ^15,16^. Here we continue to characterize those paraquat tolerance mutants by focusing on *pqt11D* in this study. We isolated *pqt11D* as a gain-of-function mutant with enhanced paraquat tolerance and identified that the *At1G01600* (*AtPQT11*) was activated by the enhancers of the T-DNA insertion, which caused paraquat resistance in the mutant. The mutant phenotype could be recapitulated by overexpressing *AtPQT11*, encoding the protein CYP86A4, a member of CYP450 super family, in the wild type Arabidopsis. Further analyses revealed that AtPQT11/CYP86A4 could convert paraquat to N-demethyl paraquat *in vitro* by the mean of N-demethylation, and demethylated paraquat was found nontoxic to Arabidopsis, clearly demonstrating paraquat degradation by a plant P450 enzyme. Our study revealed a novel molecular mechanism of paraquat resistance that has not been reported in plants. Our findings should provide a new means for developing paraquat resistance crops using *AtPQT11/CYP86A4*.

## Result

### Activated expression of *At1G01600* confers paraquat resistance in *pqt11D*

From an Arabidopsis activation-tagging library of approximately 55000 lines, we isolated a number of paraquat tolerance mutants. One of them, *pqt11D,* displayed enhanced resistance to paraquat (Fig. S1A). We found that the T-DNA with four 35S enhancers next to the right border was inserted at 1646 bp upstream the ATG codon of *At1g01600* (Fig. S1B). As a result, the gene *At1g01600* was activated (Fig, S1C), indicating that the activated expression of *At1g01600* may be responsible for observed paraquat resistance in *pqt11D.* We therefore named *At1g01600* as *AtPQT11*.

*AtPQT11* is predicted to encode CYP86A4, a cytochrome P450 enzyme with 554 amino acids. It is expressed in all tissues examined with relatively higher levels in roots, flowers and siliques (Fig. S1D), and induced by paraquat treatment for the first 3 hours, then declined (Fig. 1E).

**Figure 1.**
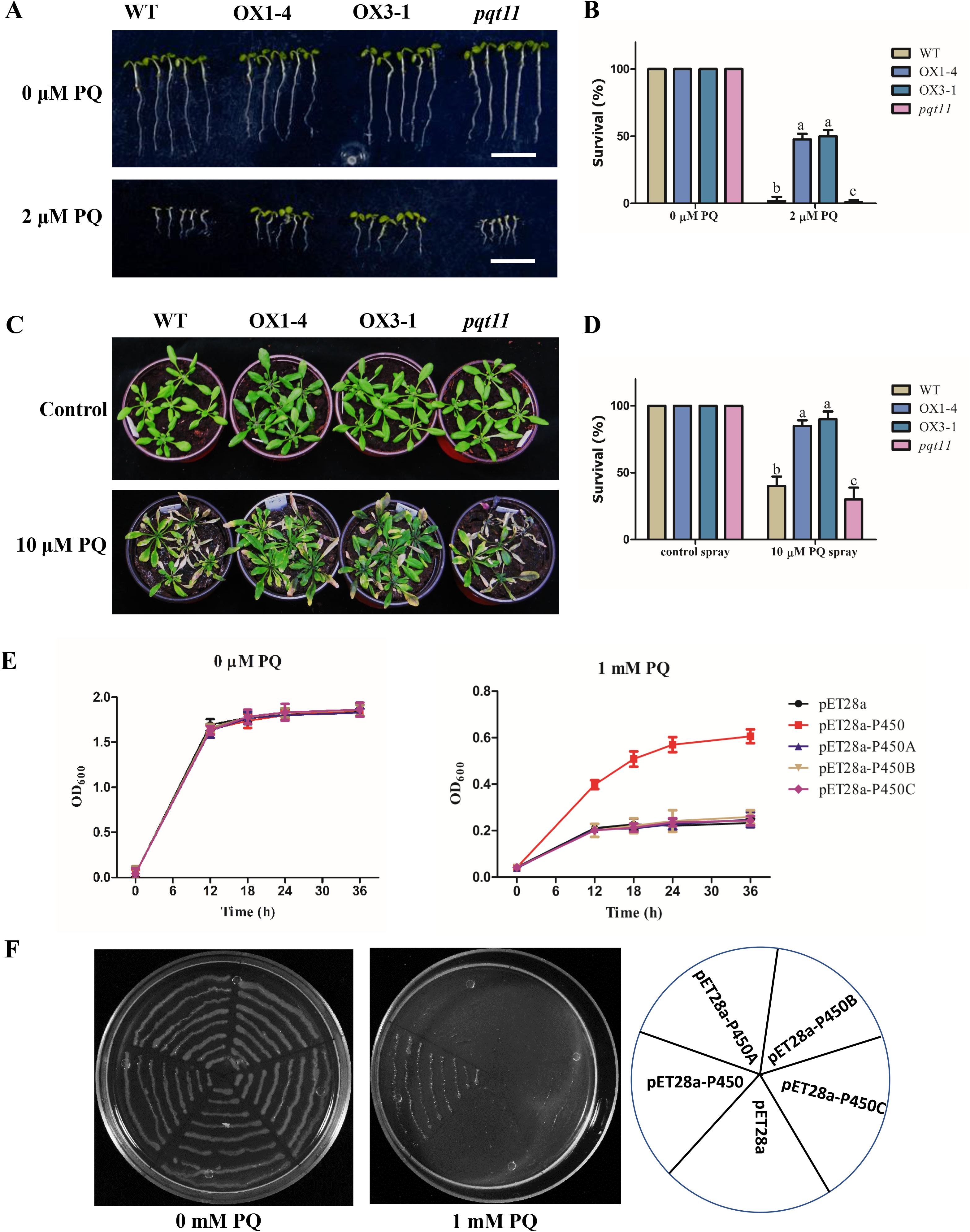
*AtPQT11* confers paraquat resistance in Arabidopsis and *E. coli* when overexpressed. (A) The seeds of wild type (WT), *AtPQT11* overexpression lines (OX1-4, OX3-1), and *pqt11* were germinated on MS medium with 0 or 2 μM paraquat (PQ) for 1 week before survival was recorded. 100 seeds were used for each genotype. 3 replicates were performed for each treatment. Bar = 1 cm. (B) Survival % in germination as in (A). Values are mean ±SD (n=3). The low case letters indicate significant differences (P < 0.05). (C) Phenotypes of wild type (WT), *AtPQT11* overexpression lines (OX1-4, OX3-1), and *pqt11* grown in the soil. When plants grew to 8-rosette-leaf stage, they were sprayed with 0 (control) or 10 μM paraquat and sprayed again 3 days later. Photos and survival % were recorded 10 days after the first spray. 40 plants were used for each genotype. 3 replicates were performed for each treatment. (D) Survival % as in (C). Values are mean ±SD (n=3). The low case letters indicate significant differences (P < 0.05). (E) Bacterial growth curves. Plasmids pET28a-AtPQT11 (pET28a-P450) with different changed residue: pET28-P450A (Gly309Ala), pET28-P450B (Cys461Ala) and pET28-P450C (Gly463Ala) were transferred into Rosetta, and cultured in LB medium with 0 or 1 mM paraquat. The growth of the cultures with initial OD_600_ adjusted to 0.1 was monitored for 36 hours. 3 replicates were performed for each treatment. Values are mean ±SD (n=3). (F) Bacterial growth on solid agar plates. Bacterial strains as in (E) were inoculated on solid LB medium with 0 or 1 mM paraquat and incubated overnight at 37°C.

### The *pqt11D* phenotype can be recapitulated by overexpressing *AtPQT11* in the wild type

To confirm whether the paraquat resistance of *pqt11D* was caused by an activated expression of *AtPQT11,* we generated two *AtPQT11* overexpression lines (OX1-4 and OX3-1) and obtained a knockout mutant *pqt11* (Salk_073078), in which the T-DNA was inserted in the first exon of *AtPQT11* from ABRC (Fig. S1B), and confirmed all the genetic materials by 3-primers PCR (Fig. S1F) and qRT-PCR analyses (Fig. S1G).

Seed germination assay in response to paraquat treatment showed that two OX lines had significantly higher survival after 7 days on MS medium with 2 μM paraquat compared with the wild type, while the knockout mutant *pqt11* became hypersensitive (Fig. 1A and B).

We also conducted paraquat resistance assay on soil-grown plants of different *AtPQT11* genotypes. When the plants grew to the stage with 8 rosette leaves, they were sprayed with 10 μM paraquat every 3 days. 10 days after the first spray, The OX lines showed a significantly higher survival than wild type control, whereas the knockout mutant exhibited sicker phenotype and lower survival (Fig. 1C and D).

Together, these results demonstrate that an elevated expression of *AtPQT11* confers paraquat resistance in Arabidopsis.

### *AtPQT11* renders *E. coli* resistant to paraquat

To further confirm *AtPQT11*-conferred paraquat resistance, we cloned *AtPQT11* cDNA into pET28a vector (pET28a-P450) and expressed in *E. coli*. Based on 3-D simulation of AtPQT11 structure (Fig. S2), we created single mutations of three residues G309A, S461A, G463A around the predicted substrate binding pocket and further expressed these three mutants in *E. coli*, respectively. In the absence of paraquat in liquid culture medium, there was no difference was observed between the bacteria expressing pET28a-P450 and the empty pET28a. In contrast, the bacteria expressing pET28a-P450 displayed a remarkable resistance to 1 mM paraquat than those harboring the empty pET28a, whereas the bacteria transformed with the constructs carrying 3 different point mutations in *AtPQT11* exhibited no paraquat resistance as the empty vector control (Fig. 1E). When grew on solid agar medium containing 1 mM paraquat, only bacteria expressing pET28a-P450 formed colony plaques (Fig. 1F). These results further confirm the role of *AtPQT11* in paraquat resistance and indicate that the amino acid residues are critical for its function.

### AtPQT11 catalyzes the conversion of paraquat to N-demethyl paraquat *in vitro*

Considering that AtPQT11 encodes a P450 protein, we reasoned AtPQT11 might catabolize paraquat to a nontoxic product. To demonstrate this, we first established an *in vitro* enzyme assay using *E. coli* expressed AtPQT11 protein, paraquat as substrate, and NADPH as electron donor as described in Materials and Methods. The consumption of NADPH provides a convenient spectrophotometric monitor at OD_340nm_ of the reaction. As shown in Fig. 2A, the consumption of NADPH was AtPQT11-dependent in the reaction and increased with incubation time. Therefore, we established the *in vitro* enzyme assay for AtPQT11 and used this assay to determine the enzyme kinetic properties (Fig. 2B-D).

**Figure 2.**
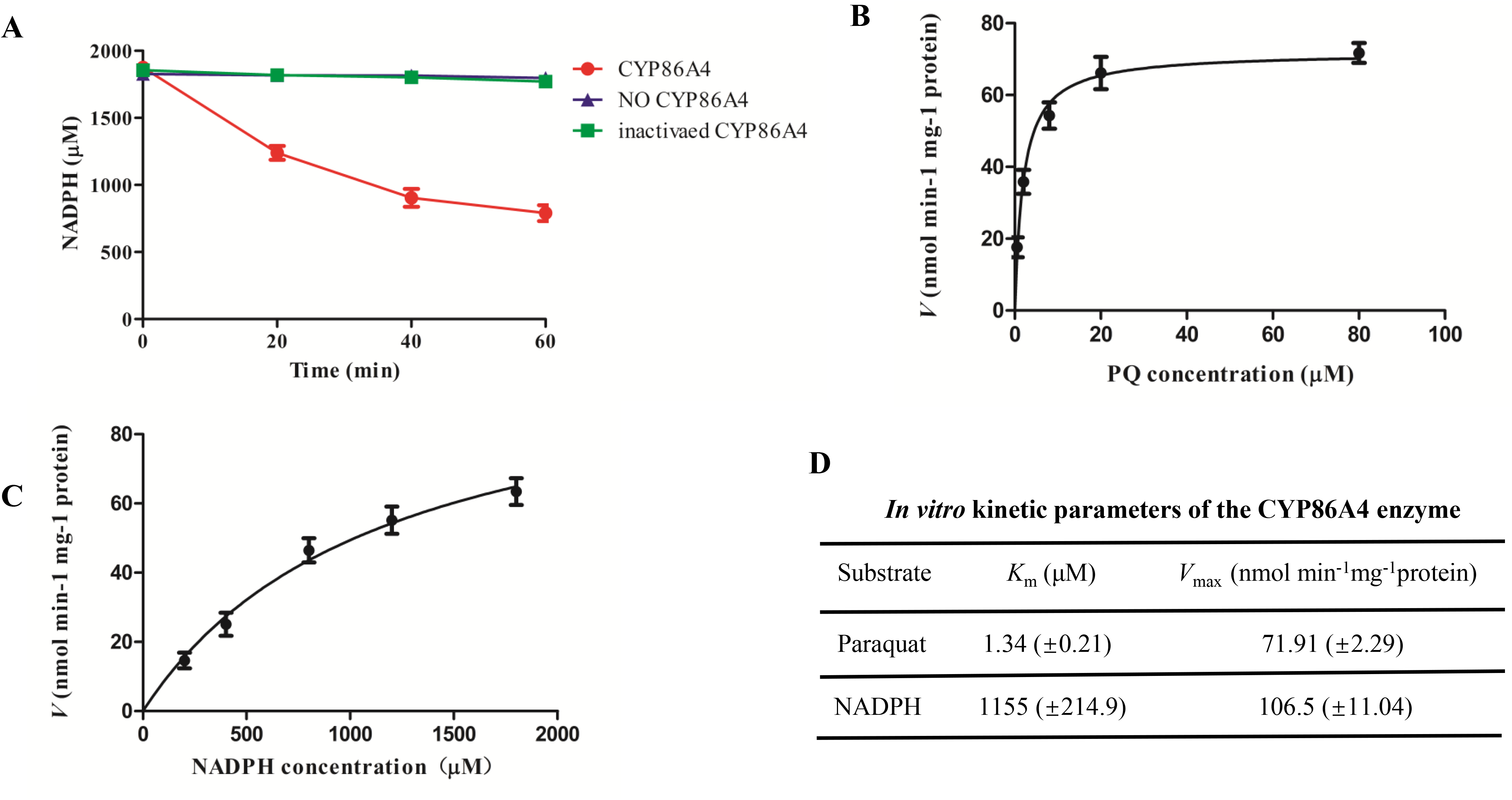
*In vitro* kinetic parameters of AtPQT11. (A) AtPQT11-dependent reaction. The enzyme assay was carried out as described in Materials and Methods with 2 controls, no AtPQt11 (CYP684A) and inactivated AtPQT11. NADPH consumption was only observed in the reaction containing AtPQT11. Values are mean ±SD (n=3). (B) Michaelis-Menton plot for paraquat. 2 mM NADPH was added to ensure it is in excess in the reaction while varying the concentration of paraquat. Values are mean ±SD (n=3). (C) Michaelis-Menton plot for NADPH. 2 mM Paraquat was added to ensure it is in excess in the reaction while varying the concentration of NADPH. Values are mean ±SD (n=3). (D) The enzyme kinetics parameters. *V_max_* and *K_m_* were obtained from (B) and (C), *K*_m_ and *V*_max_ values were estimated by using non-linear regression analysis.

To identify the product of the reaction, the paraquat metabolite of the *in vitro* reaction was analyzed by LC-MS/MS. A product peak was found at 2.7 min with m/z 171, which was AtPQT11- and NADPH-dependent (Fig. 3A-D, left panels). To confirm that this product is derived from paraquat, we used the ^2^H-labeled paraquat as the substrate, whose eight hydrogen atoms on paraquat pyridine ring were replaced by deuterium so that the m/z of paraquat changed from 186 to 194. As expected, we found the product peak with m/z 179, which was also AtPQT11- and NADPH-dependent (Fig. 3A-D, right panels). The reaction product has the same mass as N-demethyl paraquat (m/z 171).

**Figure 3.**
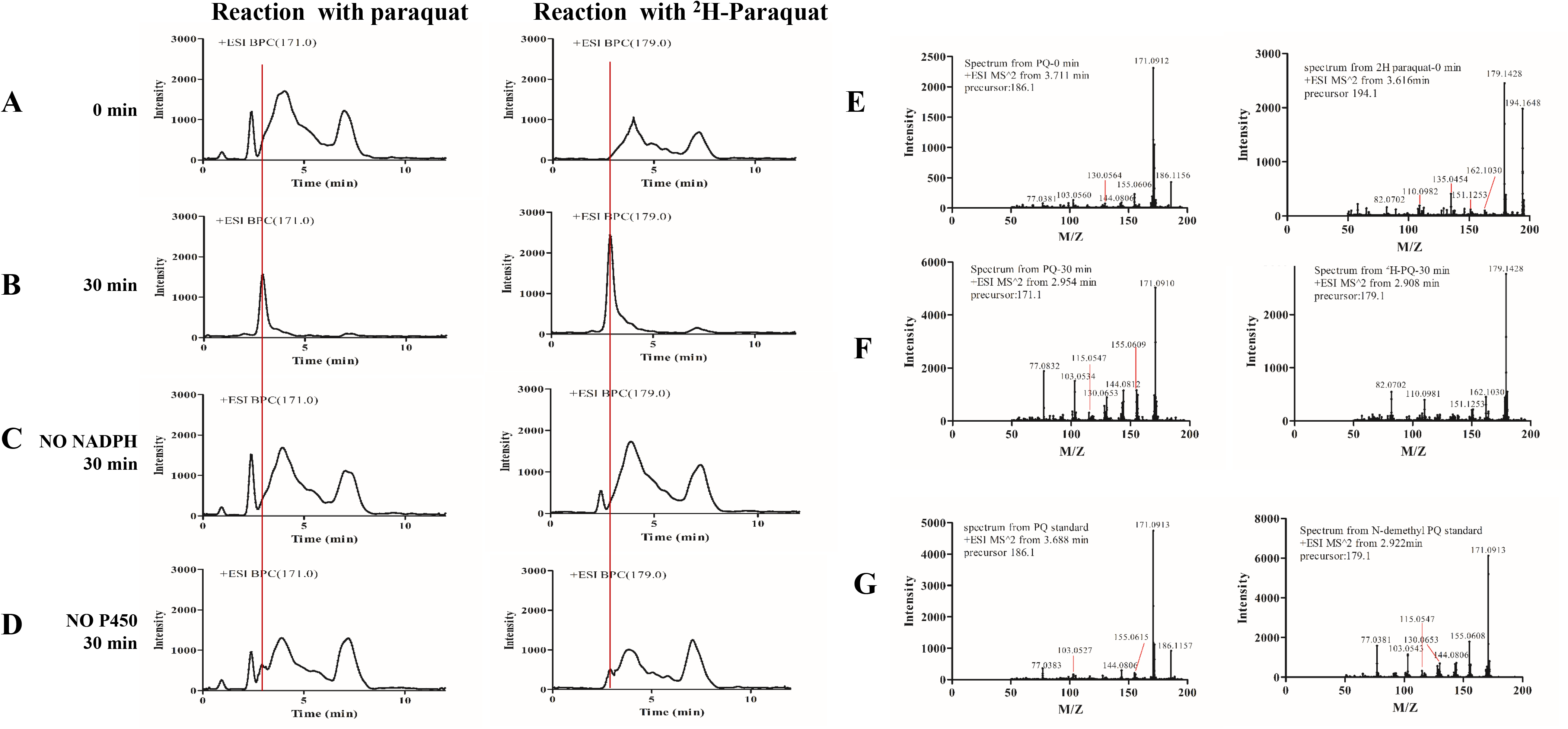
Identification of the reaction product by LC-MS/MS. (A-D) The reaction was carried out with paraquat or ^2^H-paraquat as substrate as described in Materials and Methods for 30 min. The reaction supernatant was subjected to LC-MS/MS analysis by scanning product at M/Z 171 or 179. (A) and (B) are the reaction for 0 or 30 min, (C) and (D) are the reaction for 30 min but without NADPH or P450 enzyme. (E-G) The MS/MS spectrum of substrate and product. (E) The MS/MS spectrum of paraquat when the reaction was just started. (F) The MS/MS spectrum of product when the reaction was completed (30 min). (G) The MS/MS spectrum of paraquat standard and N-demethyl paraquat standard. MS/MS spectrum of the product closely matches with that of N-demethyl paraquat (M/Z 171).

To resolve the identity of the product, we analyzed the MS/MS spectrum and found that the product showed the same fragment ions as N-demethyl paraquat standard (Fig. 3F, left panel and G, right panel), which was further confirmed by the same experiments with ^2^H-labeled paraquat (Fig. 3E and F, right panels). These results unequivocally show that the product is derived from the substrate and the product matches N-demethyl paraquat based on their fragment ion pattern (Fig. 3G). Thus, we have demonstrated that AtPQT11/ CYP86A4 functions as an N-demethylase capable of converting paraquat to N-demethyl paraquat.

### N-demethyl paraquat is nontoxic to Arabidopsis

Have identified N-demethyl paraquat as the product, we decided to find out whether N-demethyl paraquat has herbicide activity. To demonstrate this, we simply tested whether the N-demethyl paraquat is toxic to Arabidopsis. Seed germination assay in the presence of N-demethyl paraquat showed that two OX lines, WT and knockout mutant *pqt11* all survived and grew well on MS medium with 2 μM N-demethyl paraquat or 2 μM bipyridine (double demethylated paraquat) compared with paraquat (Fig. 4A and B). This result clear shows that N-demethyl paraquat and bipyridine are nontoxic to Arabidopsis. Therefore, we have uncovered the molecular mechanism underlying the paraquat resistance of *pqt11D*.

**Figure 4.**
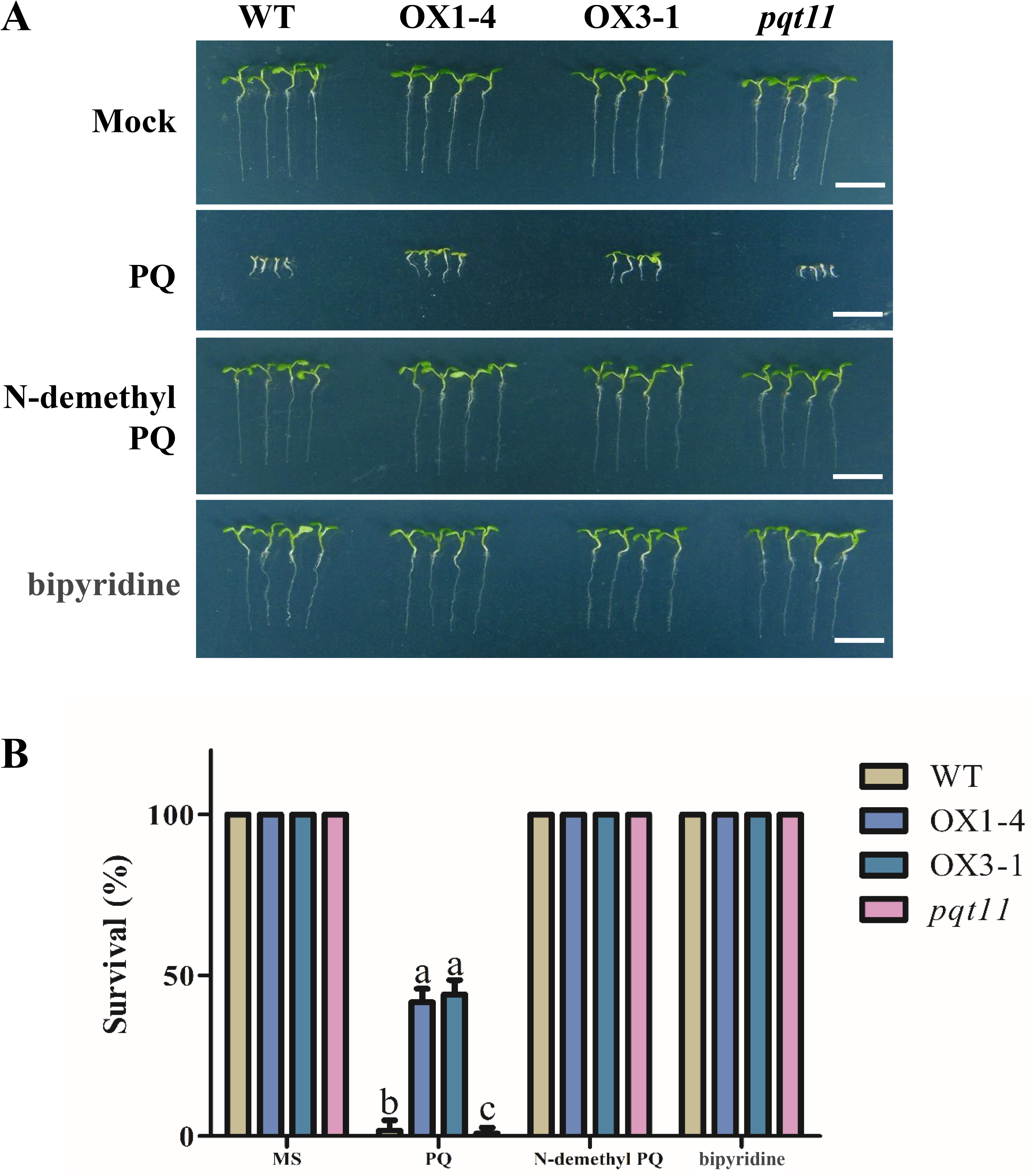
N-demethyl paraquat is nontoxic to Arabidopsis. (A) Germination assay. The seeds of wild type (WT), *AtPQT11* overexpression lines (OX1-4, OX3-1), and *pqt11* were germinated on MS medium with 0 (Mock), 2 μM paraquat (PQ), 2 μM N-demethyl paraquat (N-methyl PQ), or 2 μM bipyridine, respectively for 1 week before survival was recorded. 100 seeds were used for each genotype. 3 replicates were performed for each treatment. Bar = 1 cm. (B) Survival % in germination as in (A). Values are mean ±SD (n=3). The low case letters indicate significant differences (P < 0.05).

## Discussion

In this study we reported the paraquat resistance mutant *pqt11D* and the underlying molecular mechanism. The phenotype of the mutant was caused by an activated expression of *AtPQT11*, which was confirmed by recapitulation analyses in Arabidopsis wild type as well as in *E. coli* (Fig. 1 and Fig. S1A-C). The paraquat hypersensitive phenotype of the *AtPQT11* knockout mutant also supports that *AtPQT11* is responsible for the observed paraquat resistance (Fig. 1 and 4). Further analyses revealed the AtPQT11 catalyzed biochemical reaction converting paraquat to single demethylated product (Fig. 2 and 3), and we also found that N-demethyl paraquat is nontoxic to Arabidopsis (Fig. 4).

CYP450 enzymes is widely found in biosynthesis or catabolic processes through the whole life in all living organisms ^22^. Several members of CYP450 family have been reported to detoxify herbicides ^23,24^. As a xenobiotic, paraquat is not a natural substrate of the CYP450 enzyme AtPQT11. The natural substrates of AtPQT11 remain unknown at present. AtPQT11 can catalyze the conversion of paraquat to N-demethyl paraquat perhaps because paraquat may have similar structural resemblance to its natural substrates.

Since paraquat is a widely used herbicide in agriculture, paraquat resistant weeds have emerged, which is a potential threat to agriculture. Generally, plants could gain paraquat resistance in three ways. First, paraquat absorption and transport are limited, thus preventing paraquat from reaching chloroplasts. Second, an enhanced intracellular antioxidant capacity would help plants to scavenge ROS more efficiently. Third, paraquat may be degraded to nontoxic metabolites. Plants may have these strategies in single or in combination to produce paraquat resistance ^25^. There are many reports for the first two strategies ^4,10,13,15,16^. However, paraquat degradation by plants has not been reported thus far. Our finding with *pqt11D* fills up the gap for the third strategy.

In conclusion, our findings with the paraquat resistant mutant *pqt11D* unequivocally demonstrate that AtPQT11, a member of the P450 super family detoxifies paraquat and confers paraquat resistance in Arabidopsis when overexpressed, therefore providing a much needed means to engineering paraquat resistance crops.

## Methods and materials

### Plant material and growth conditions

Arabidopsis Col-0 was used as wild type in this study. We also used Col-0 as the genetic background for all the mutants and transgenic plants. Salk_073078 was ordered from the Arabidopsis Biological Resource Center (ABRC). Seeds were sterilized in 10% bleach for 10 minutes and washed with sterile water 5 times. The washed seeds were kept in the dark at 4°C for 2 days before germination on MS medium at 22°C under 14 h light/10 h dark.

### Construction of AtPQT11 overexpression lines

In order to get overexpression lines, we used forward primer 5’-GGGGACAAGTTTGTACAAAAAAGCAGGCT ATGGAAATATCCAATGCCATGC-3 and reverse primer 5’-GGGGACCACTTTGTACAAGAAAGCTGGGT TTAAACCACTGCAACTCCCGTA-3’ to obtain the CDS of CYP86A4. By means of GATEWAY system ^26^, the CDS was cloned into vector pCB2004 via the shuttle vector of pDONR207.

The constructed plasmid was transformed into *Agrobacterium tumefaciens* C58C1 by electroporation. The floral-dip method was used to transfer the construct into plants ^27^. The positive lines were screened as glufosinate resistant plants and homozygous lines were obtained from F2 population.

### RNA extraction and qRT-PCR

We extracted the RNA from one-week-old seedlings by Trizol (Invitrogen, Carlsbad, California, USA) and reverse transcribed RNA to cDNA by TransScript Kit (TaKaRa). The expression of CYP86A4 was detected by RT-PCR or quantitative RT-PCR using forward primer 5’-CCCCAAGGGTTTCACTGAATTC-3’ and reverse primer 5’-AAGTAAATGCGAAGCCTGCTTG −3’ and Applied Biosystem Step One real-time PCR system or TaKaRa SYBR Premix Ex Taq II reagent kit. *UBQ5* was used as the internal reference.

### Identification of the *AtPQT11* knockout mutant

3 primers including forward primer 5’- TTCACCACATACAGCTGCATC −3’、 reverse primer 5’- AAATGTTGTCGAATGTGAGCC-3’ and intermediate primer LBb1.3 5’-ATTTTGCCGATTTCGGAAC-3’ were used to identify homozygotes of SALK_073078. The homozygotes showed only one band around 750 bp while the wild type showed one band around 1000 bp. RT-PCR was performed to confirm null expression of the *At1g01600* in SALK_073078 by using forward primer 5’- ATGGAAATATCCAATGCCATGC-3’ and reverse primer 5’- TTAAACCACTGCAACTCCCGTA-3’. *TUBULIN* was used as the internal control.

### Seed germination assay

Sterilized seeds for each line including WT, knock out mutant SALK_073078 (*pqt11*/*cyp86a4*) and two overexpression lines OX1-4 and OX3-1 were treated at 4°C in dark condition for 2 days. Then we put these seeds on MS medium with 2 μM paraquat, MS medium was used as control. The plates were placed at 22°C under 14 h light/10 h dark cycles. We considered the appearance of two green cotyledons as successful germination, 1 week later we recorded germination %.

### Paraquat resistance assay on soil-grown plants

One-week-old seedlings grown on MS medium were transferred to soil. We planted 5 seedlings per pot, grew to the stage with 8 rosette leaves, and sprayed with 10 μM paraquat and sprayed again 3 days later. Photos and survival rate were recorded 10 days after the first spray.

### 3-D model of At PQT11 structure

The 3-D structure model of CYP86A4 was produced by Phyre2 and paraquat was docked into the structure model. Residues within the distance of 4 Å around paraquat are colored in red. Conservative residues likely responsible for the binding of paraquat are labeled and chosen as mutation sites.

### Site-directed mutagenesis of *AtPQT11*

Specific primers for site-directed mutagenesis were designed, for changing 309Gly to Ala, 461Cys to Ala, 463Gly to Ala as listed in Table S1. pET28-CYP86A4 or pCB2004-CYP86A4 were set as template and the PCR procedure was set for 20 cycles. After the PCR products were recycled, 1 μl DpnⅠwas added to digest these products. Then transferred these products to DH5α and extracted plasmids from overnight culture.

### Paraquat resistance assay in *E. coli*

We used Rosetta or BL21 as host, transferred the constructed pET28-P450 and the three plasmids with point mutation: pET28-P450A (Gly309Ala), pET28-P450B (Cys461Ala) and pET28-P450C (Gly463Ala) into the host bacteria, and empty vector pET28a as control. Single colonies grown on the plate were picked and cultured overnight. Then the overnight culture was diluted into fresh LB medium with a starting OD_600_ value of 0.1, and paraquat was added to make its final concentration of 1 mM. Another set of culture with 0 mM paraquat was used as control. The OD_600_ value of the culture was monitored at regular intervals for 36 hours. Each culture was triplicated and the growth curves were obtained. When tested on solid LB plate with 1 mM paraquat, the overnight liquid bacteria culture was inoculated on the plate with 0 or 1 mM paraquat and incubated overnight at 37°C.

### Bacterial expression of AtPQT11

The CDS of *AtPQT11* was ligated to the vector pET28a by T4 ligase. The constructed plasmid was transformed into Rosetta strain with kanamycin selection. A clone was picked from medium and was added with liquid LB, when the OD_600_ of medium reached 0.6, IPTG was added with a final concentration of 0.3 mM. The culture was shaken in 16°C with the speed of 120 rpm for 12-16 h. The culture was centrifuged for 10 min at 4500 rpm to harvest bacteria, then the bacterial pellet was resuspended with protein isolation buffer (20 mM Tris-Cl pH7.5, 0.2 M NaCl, 5% glycerin, 1 mM EDTA, 1 mM PMSF) + 1% Dodecyl-beta-D-maltoside (m/v), and ultrasonicated to break bacterial cell walls. The cell lysate was rotated at 4°C for 2h and then centrifuged at 45000 rpm for 30 min. The supernatant was collected for AtPQT11 protein purification using Ni-IDA-sefinose resin kit.

### *In vitro* enzyme assay of AtPQT11

The reaction mixtures contained 0.1-10 μM paraquat as substrate, 50-100 μg AtPQT11/ CYP86A4 protein, 1 mM NADPH as electron donor, 25 mM phosphate buffer (pH7.0) in a total volume of 50 μl. The reaction was initiated by adding NADPH. The reaction mixtures without NADPH or AtPQT11/CYP86A4 were set as controls. The mixtures were incubated for 30 min at 27°C and terminated by adding 50 μl acetonitrile. To determine the enzyme kinetic properties, the reaction was spectrophotometrically monitored at 340 nm for NADPH consumption. To identify the reaction product, the mixtures was centrifuged at 14000g for 20 min, and the supernatant was analyzed by HPLC-MS/MS as described below.

### HPLC-MS/MS

HPLC-MS/MS analysis was performed using Agilent Q-TOF-LC/MS 6545 device (Agilent Technologies, Palo Alto, CA, USA). A XBridge BEH HILIC 2.5μm Column (2.1*100 mm) from Waters was used for chromatographic separation. The mobile phase was a mixture of solvent A (10 mM ammonium acetate added with 0.1% V/V formic acid) and solvent B (10 mM ammonium acetate dissolved in 95% acetonitrile) and was delivered at a flow rate of 0.3 mL/min. The column temperature was set at 40°C and the injection volume was 2 μl. The sample was followed the gradient: 90% B (0 min), 10% B (3 min), 1% B (5 min), 90% B (5 min), with a total run time of 12 min.

Mass acquisition was performed in the positive ionization mode at a fragmentation voltage of 175 V. The following parameters were used: drying gas flow, 8 L/min; temperature, 325 °C; sheath gas flow, 11 L/min; temperature, 350 °C; nebulizer pressure, 45 psi; and capillary voltage, 3500 V. The collision energy was set at 15–40 V, and the mass range was recorded from m/z 50–1700. The acquired MS and MS/MS data files were analyzed using Agilent MassHunter Qualitative Analysis B.07.00.

### Toxicity assay of N-demethyl paraquat

Seeds of different genotypes were washed and treated at 4°C in dark condition for 2 days, then plated on MS medium with 2 μM N-demethyl paraquat, 2 μM bipyridine, 2 μM paraquat, respectively. MS medium was used as mock. Germination and survival was recorded after 1 week.

### Statistical analyses

Statistical significance was evaluated at the 0.05 probability level using Student’s t-test.

## Supplemental information

Figure S1. Paraquat resistance phenotype of *pqt11D* and confirmation of

*AtPQT11* expression in *pqt11*, *pqt11D*, and overexpression lines.

Fig. S2. 3-D model of AtPQT11 structure.

Table S1. Primers used in this study.

## Acknowledgements

This work was supported by grants from the National Natural Science Foundation of China (grant no. 31770273). We thank ABRC for providing Arabidopsis mutant seeds.

## Author’s Contributions

CBX and YJH designed the experiments. YJH. performed the major experiments. YPH isolated *pqt11D* and identified the T-DNA insertion site. JQX, WTH, AMM, XQQ, YXC, participated in experiments and data analyses. ZPF, YFC, YiPH, LPG contributed to LC-MS/MS analyses. YJH wrote the manuscript. CBX and LPG revised the manuscript and supervised the project.

