## Supplementary material for "AtPQT11, a P450 enzyme, detoxifies paraquat via N-demethylation": Supplemetal information: Supplemental info.pdf

**Supplemental information for “AtPQT11, a P450 enzyme, detoxifies paraquat by N-demethylation” by Huang et al.**

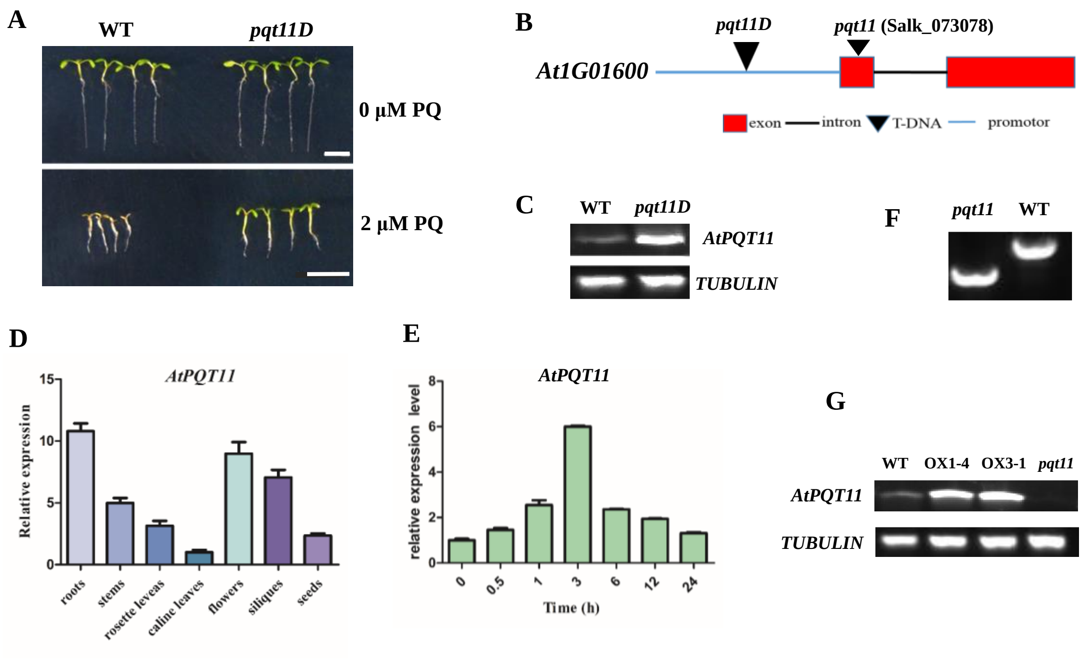


**Fig. S1. Paraquat resistance phenotype of *pqt11D* and confirmation of *AtPQT11* expression in *pqt11*, *pqt11D*, and overexpression lines.**

A. Paraquat resistance phenotype of *pqt11D* vs. wild type. The mutant *pqt11D* displayed enhanced paraquat resistance on MS medium containing 2 μM paraquat (PQ) while the wild type was inhibited. Bar=1cm.

B. Schematic representation of *AtPQT11* gene and location of T-DNA insert sites (indicated by triangle) in *pqt11* and *pqt11D*.

C. RT-PCR analysis of *AtPQT11* in *pqt11D* vs. wild type (WT). The expression of *AtPQT11* was significantly activated in *pqt11D* compared with that of wild type. *TUBULIN* was used a reference.

D. Quantitative RT-PCR analysis of *AtPQT11* expression in different tissues.

E. Quantitative RT-PCR analysis of *AtPQT11* expression in response to paraquat treatment.

F. Verification of Salk_073078 /*pqt11* at genomic level by the standard 3-primers procedure.

G. RT-PCR analysis of *AtPQT11* in wild type (WT), two OX lines (OX1-4, OX3-1) and *pqt11*. *TUBULIN* was used a reference.

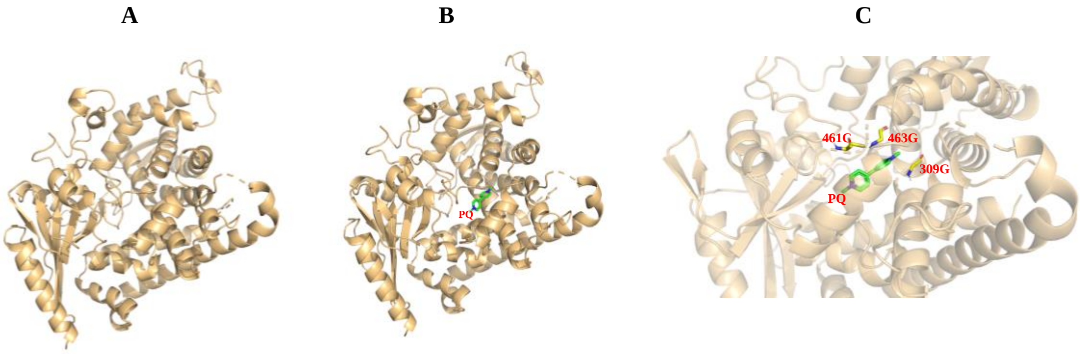


**Fig. S2. 3-D model of AtPQT11 structure.**

A. The 3-D structure model of CYP86A4 produced by Phyre2

B. Docking of paraquat into the structure model.

C. Three conservative residues (309G、461C、463G), which are within the distance of 4 Å around paraquat and likely responsible for the binding of paraquat, are shown as sticks.

**Table S1. Primers used in this study.**

| Name | Primer Sequence | vector |
|---|---|---|
| PQT11-CDS | P1:GGGGACAAGTTTGTACAAAAAAGCAGGCT ATGGAAATATCCAATGCCATGC | pCB2004 (for PQT11 gene Overexpression) |
| | P2:GGGGACCACTTTGTACAAGAAAGCTGGGT TTAAACCACTGCAACTCCCGTA | |
| SALK_073078 | P1:TTCACCACATACAGCTGCATC | |
| | P2: AAATGTTGTCGAATGTGAGCC | |
| | LBb1.3: ATTTTGCCGATTTCGGAAC | |
| PQT11-qPCR | P1:CCCCAAGGGTTTCACTGAATTC | |
| | P2: AAGTAAATGCGAAGCCTGCTTG | |
| β －tubulin8 | P1: CTTAAGCTCACCACTCCAAGCT | |
| | P2: GCACTTCCACTTCGTCTTCTTC | |

| | | |
|---|---|---|
| Ubiquitin5 | P1: AGAAGATCAAGCACAAGCAT | |
| | P2: CAGATCAAGCTTCAACTCCT | |
| pET28a-PQT11-Nde 1<br>pET28a-PQT11-Sma 1 | P1:GGAATTCCATATG ATGGAAATATCCAATGCCATGC | pET28a (for PQT11/CYP86A4 protein expression) |
| | P2:TCCCCCGGG TTAAACCACTGCAACTCCCGTA | |
| PQT11-309A | P1:AACTTTATCCTAGCTGCACGTGACACGT | pET28a (for PQT11/CYP86A4 protein point mutation) |
| | P2:GCACCGCGAATTGAAATAGGATCGACG | |
| PQT11-461A | P1:AATGCTGGACCAAGGATCGCATTGGGGAAA GATCTGGCG | |
| | P2:TTACGACCTGGTTCCTAGCGGAACCCCTTTC TAGACCGC | |
| PQT11-463A | P1:CCAAGGATCTGCTTGGCGAAAGATCTG | |
| | P2:GTTACGACCTGGTTCCTAGACGAACCG | |
